# An actuated neural probe architecture for reducing gliosis-induced recording degradation

**DOI:** 10.1101/380006

**Authors:** Travis L. Massey, Leane S. Kuo, Jiang Lan Fan, Michel M. Maharbiz

**Affiliations:** Dept. Electrical Engineering and Computer Sciences, University of California, Berkeley; Dept. Bioengineering, University of California, Berkeley; UC Berkeley/UCSF joint graduate program in bioengineering; Helen Wills Neuroscience Institute, University of California, Berkeley; Center for Neural Engineering and Prostheses, University of California, Berkeley/San Francisco; Chan Zuckerberg Biohub, San Francisco, CA 94158

## Abstract

Glial encapsulation of chronically implanted neural probes inhibits recording and stimulation, and this signal loss is a significant factor limiting the clinical viability of most neural implant topologies for decades-long implantation. We demonstrate a mechanical proof of concept for silicon shank-style neural probes intended to minimize gliosis near the recording sites. Compliant whiskers on the edges of the probe fold inward to minimize tissue damage during insertion. Once implanted to the target depth and retracted slightly, these whiskers splay outward. The splayed tips, on which recording sites could be patterned, extend beyond the typical 50-100 micron radius of a glial scar. The whiskers are micron-scale to minimize or avoid glial scarring. Electrically inactive devices with whiskers of varying widths and curvature were designed and monolithically fabricated from a five-micron silicon-on-insulator (SOI) wafer, and their mechanical functionality was demonstrated in a 0.6% agar brain phantom. Deflection was plotted versus deflection speed, and those that were most compliant actuated successfully. This probe requires no preparation for use beyond what is typical for a shank-style silicon probe.

## I. Introduction

Intracortical electrophysiological neural recording is a ubiquitous technique for detecting single-unit activity in the central nervous system, and recent decades have seen a plethora of devices developed toward this end [1–3]. Popular among these are shank-style silicon probes, in which electrodes for recording or stimulation are arranged along the body of a rigid silicon probe [4–9]; such probes can be microfabricated with relatively high throughput and are amenable to CMOS integration. When these probes are inserted into the brain, they initiate an adverse biological response that notably includes the formation of a 50-100 μm glial scar around the implanted foreign body [10]. This scar displaces healthy tissue, degrading the quality of recorded units and increasing signal amplitudes required to electrically stimulate nearby neurons [11–13]. Reducing the crosssectional dimensions of the implant to single-digit-micron scales can reduce or eliminate gliosis [12, 14], but this fine structure limits the probe length to no more than a few millimeters before it buckles upon attempted insertion into the tissue [15, 16]. Seymour and Kipke demonstrated reduced encapsulation around a sparse parylene-C scaffold extending off of a probe body [12], but the vasculature damage caused during insertion initiates a local immune response near the final position of the recording sites [10].

A common feature of nearly all neural recording arrays is that they are static. Even those fabricated with compliant materials move only passively in response to their surroundings (i.e. brain micromotion). Dynamic probes, in contrast, incorporate traditional MEMS principles to actively move. This allows one to position a recording or stimulation site on a fine structural member distal to the probe body and out of the radius of the glial scar. To date, there are very few examples of such devices in the literature. Egert and Najafi demonstrated probes with spring-loaded recording sites that would deploy upon timed dissolution of a polymer post-implantation [17]. Xie *et al*. demonstrated pre-strained bimetallic strip electrodes that were frozen in liquid nitrogen in a flat position, implanted while frozen, and curled outward away from the body of the probe as they thawed [18]. In both cases, deployment was timed by some physical mechanism and required nontrivial preparation before implantation. While Egert showed that the deployment timing could be controlled by the choice of resorbable polymer adhesive [19], the surgeon is limited to this fixed deployment time regardless of case-by-case delays that may arise during implantation.

We present mechanical proof of concept for a simple actuated probe monolithically microfabricated in silicon. This design requires no preparation before implantation beyond what is typical for static shank-style probes and can be deployed at will following implantation. Compliant whiskers extending from the probe body fold against the body during implantation, minimizing vasculature damage, and splay outward when the probe is retracted slightly at the target depth. Recording sites, if patterned at the tips of the deployed whiskers, would extend beyond the reach of the glial scar surrounding the probe body, and the whiskers are sufficiently fine that no glial encapsulation is expected in the vicinity of the recording sites [12, 14, 15, 20, 21]. The following sections detail the design of these devices, hereafter referred to as “splaying probes,” followed by the microfabrication and testing procedures, the results of insertions in a agar brain phantoms, and preliminary results in porcine brain tissue.

## II. Probe Design

### A. Conceptual design

The probes were designed to have the archetypal shape of a shank-style silicon probe with a wide, thin body tapering to a 50° point. Whiskers on each side of the probe body extend outward, parallel to the probe body at the anchor point and curving outward with a circular profile. Each whisker subtends an angle of *π*/4 radians, such that when the whisker is fully folded inward by *π*/4 radians it rests flat against the body of the probe, and when splayed outward by *π*/4 radians the tip is perpendicular to the probe body. The circular profile was chosen in order that the stress in the whisker would be evenly distributed when folded flat into the body. The radius of this circle is hereafter referred to as the “arc radius,” *r*.

### B. Quantitative design

Once the whisker shape was established, dimensions were determined according to two key criteria. First, the insertion force must be sufficient to fully fold the whiskers in toward the body of the probe. Because the insertion force is not well defined and a function of the probe geometry, Roark’s canonical beam bending equations (1) were used to determine the minimum arc radius such that the beam would deflect by the full *π*/4 radians for a broad range of forces (Figure 1a) [22]. Curves are plotted for 3, 4, 5, and 6μm beam widths. These values are sufficiently small to minimize the adverse biological response when implanted, but sufficiently large for ease of fabrication, handling, and future electrode patterning. Note that the forces plotted are the equivalent drag force as a point load at the whisker’s tip, not a measure of the total distributed insertion force.

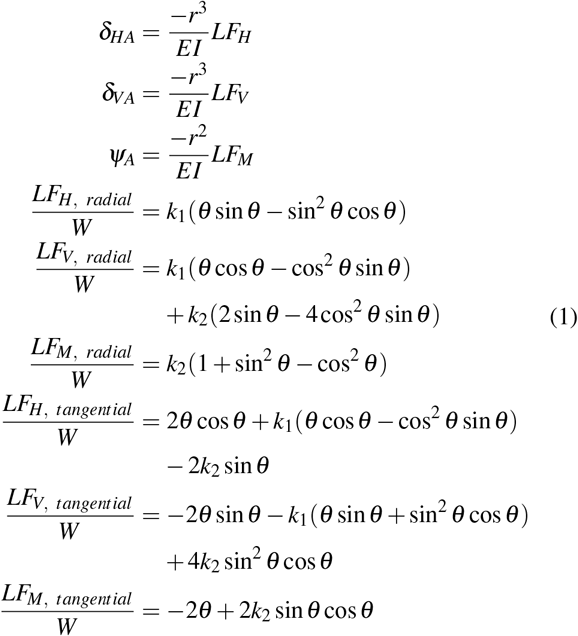

**Fig. 1.**
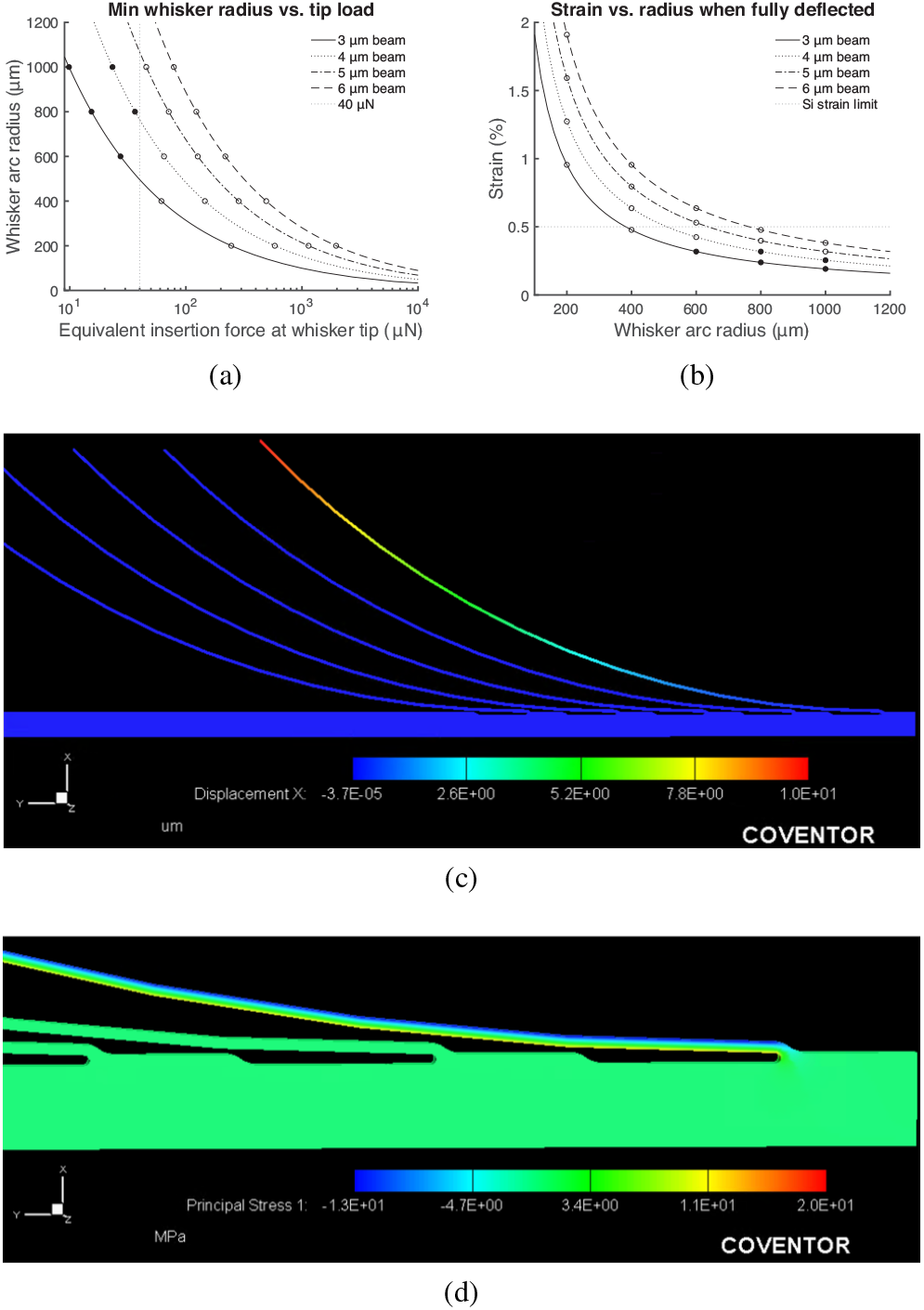
(a) Plot of the minimum whisker arc radius to fully deflect inward (*π*/4 radians) in response to a point load applied at the whisker’s tip. The region above each curve corresponds to the the valid design space for that beam width. Circles indicate those devices fabricated; filled circles indicate devices that folded fully during insertion, and open circles indicate those that did not. The gray vertical dotted line suggests an approximate equivalent insertion force, discussed further in Section IV. (b) Plot of strain versus whisker arc radius, with a horizontal line indicating the approximate yield strain of single-crystal silicon micromachined by deep reactive ion etching (DRIE). (c) Simulated displacement of a whisker (*w* =3μm, *r* =1000μm) to verify calculated spring constant. (d) Simulated stress distribution in deflected whisker.

In the above equations, *LF* are the load terms for the corresponding horizontal, vertical, and rotational (moment) deflections for radially and tangentially loaded curved beams. These equations apply for beams with one end free and the other fixed, and the point load applied at the tip of the free end. *W* is the applied load, and ***θ*** is the half of the beam’s arc angle (*π*/8). Correction factors *k*_1_ and *k*_1_ are within 0.11.0% of 1 because the beam has a large radius relative to its width. ***δ*** and ***ψ*** are linear and rotational deflections at the tip. The direction of the deflections ***δ***_*HA*_ and ***δ***_*VA*_ are tangential and perpendicular to the midpoint of the beam, respectively. To find the horizontal and vertical deflections in the frame of reference of the probe, we applied a rotation matrix to the calculated ***δ***_*HA*_ and ***δ***_*VA*_.

The second design criterion states that the beam should not break when deflected; that is, the strain at the beam’s surface (***ε***) under maximum deflection should not exceed the yield strain of the silicon (2). The exact yield strain is process dependent, but in this process it is typically ≥0.5%. *l* and *w* are the length and width of the beam, respectively. Strain as a function of beam radius is plotted in Figure 1b.

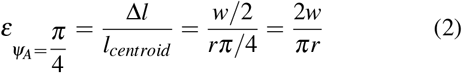

To accommodate uncertainty in the approximations of insertion force and yield strain, a range of beams spanning three orders of magnitude of relative stiffness were fabricated; *w* = 3, 4, 5, and 6μm, and *r* = 200, 400, 600, 800, and 1000 μm. Longer or thinner whiskers were impractical to fabricate and handle, and shorter or thicker whiskers violated one of the aforementioned constraints. The whiskers’ spring constants in response to a point load applied at the tip of the beam parallel to the probe body are listed in Table I (parallel to loading direction) and Table II (perpendicular to loading direction), and the rotational compliance values are provided in Table III.

**TABLE I.** Calculated curved beam spring constants (*N/m*) in loading direction for point load applied at tip, parallel to probe body

| Beam width | $r=200\ \mu\text{m}$ | $r=400\ \mu\text{m}$ | $r=600\ \mu\text{m}$ | $r=800\ \mu\text{m}$ | $r=1000\ \mu\text{m}$ |
| --- | --- | --- | --- | --- | --- |
| 3 $\mu\text{m}$ | 6.7105 | 0.8391 | 0.2486 | 0.1049 | 0.0537 |
| 4 $\mu\text{m}$ | 15.9002 | 1.9888 | 0.5894 | 0.2486 | 0.1273 |
| 5 $\mu\text{m}$ | 31.0397 | 3.8840 | 1.1510 | 0.4856 | 0.2486 |
| 6 $\mu\text{m}$ | 53.6041 | 6.7105 | 1.9888 | 0.8391 | 0.4296 |

**TABLE II.** Calculated curved beam spring constants (*N/m*) perpendicular to the loading direction for point load applied at tip, parallel to probe body

| Beam width | $r=200\ \mu\text{m}$ | $r=400\ \mu\text{m}$ | $r=600\ \mu\text{m}$ | $r=800\ \mu\text{m}$ | $r=1000\ \mu\text{m}$ |
| --- | --- | --- | --- | --- | --- |
| 3 $\mu\text{m}$ | 3.6893 | 0.4612 | 0.1367 | 0.0577 | 0.0295 |
| 4 $\mu\text{m}$ | 8.7438 | 1.0932 | 0.3239 | 0.1367 | 0.0700 |
| 5 $\mu\text{m}$ | 17.0751 | 2.1351 | 0.6327 | 0.2669 | 0.1367 |
| 6 $\mu\text{m}$ | 29.4999 | 3.6893 | 1.0932 | 0.4612 | 0.2361 |

**TABLE III.**
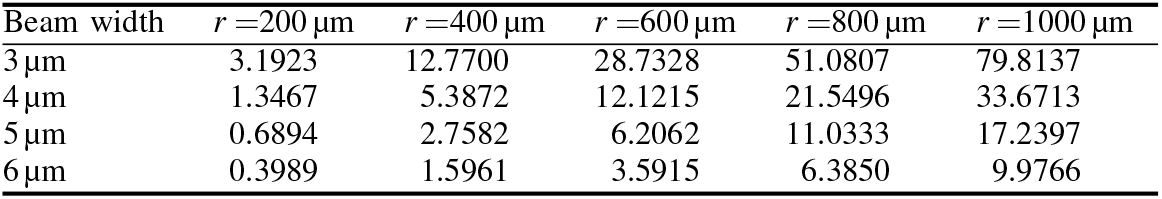
Calculated curved beam rotational compliance (*rad/μN*) for point load applied at tip, parallel to probe body

| Beam width | $r=200\ \mu\text{m}$ | $r=400\ \mu\text{m}$ | $r=600\ \mu\text{m}$ | $r=800\ \mu\text{m}$ | $r=1000\ \mu\text{m}$ |
| --- | --- | --- | --- | --- | --- |
| 3 $\mu\text{m}$ | 3.1923 | 12.7700 | 28.7328 | 51.0807 | 79.8137 |
| 4 $\mu\text{m}$ | 1.3467 | 5.3872 | 12.1215 | 21.5496 | 33.6713 |
| 5 $\mu\text{m}$ | 0.6894 | 2.7582 | 6.2062 | 11.0333 | 17.2397 |
| 6 $\mu\text{m}$ | 0.3989 | 1.5961 | 3.5915 | 6.3850 | 9.9766 |

To verify these hand calculations, we simulated the deflection of one whisker with *w* =3 μm and *r* =1000 μm in CoventorWare (Coventor, Inc, Cary, NC). The spring constants matched the hand-calculated values within 6%. A plot of the stress distribution also confirmed that the stress was distributed throughout the whisker, though there was a slight concentration near the base. Images from these simulations are provided in Figure 1c-d.

## III. Microfabrication and Testing Procedure

### A. Device layout and microfabricaiion

The layout was procedurally generated in MATLAB (MathWorks, Natick, MA) and AutoCAD (Autodesk, San Rafael, CA) using custom scripts. Whisker anchor points were rounded and filleted to minimize stress concentrations, and whiskers were aligned at the base such that the probe did not widen as whiskers were added. Figure 2a-b shows the probe layout and a close-up of the anchor points.

**Fig. 2.**
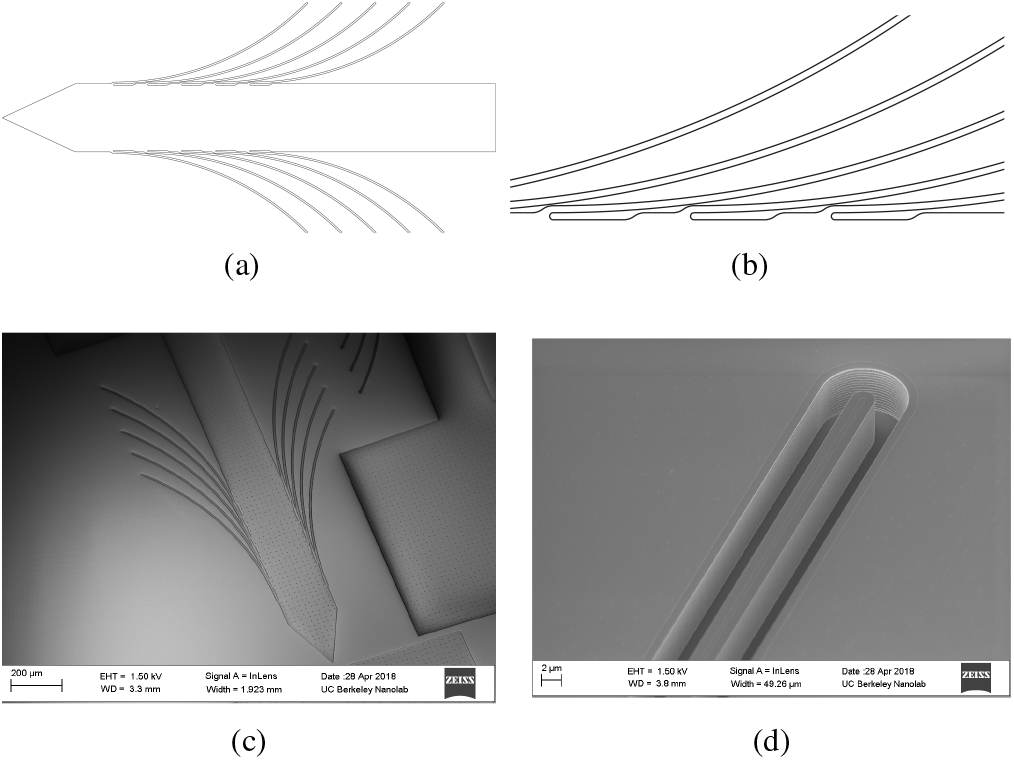
(a) Layout of a full probe, without handle. Shank width is 200 μm. (b) Close-up of layout showing the base of the 5μm-wide whiskers. (c) SEM of a representative probe, etched but not released. (d) Close-up SEM of a 3 μm-wide, 5 μm-tall whisker surrounded by a 3 μm-wide etched trench.

Probes were microfabricated from a 5 μm device layer silicon-on-insulator wafer using standard cleanroom processes. Photoresist was patterend in the outline of the device, and a trench was etched by DRIE, yielding structures as seen in Figure 2c-d. After stripping the photoresist, the devices were released using vapor-phase hydrofluoric acid. Etch holes were regularly patterned along the probe body and handle to aid in the release step. The holes had small random offsets to reduce the risk of creating a weak point for fracture along the silicon’s crystal plane. Following release, the devices could be conveniently manipulated by a 1000 by 850 μm handle (not shown in Figure 2a; visible on right edge of 2c).

### B. Insertion tests in agar brain phantom

The mechanical operation of the probes was tested in 0.6 weight percent agar, the standard mechanical phantom for cortical matter (specifically excluding the meninges) [23–25]. The agar (9002-18-0, Millipore-Sigma, Darmstadt, Germany) was prepared on a hot plate by standard techniques and allowed to fully gel in a 10 mm cuvette before use. A thin film of deionized water was applied to the top surface of the agar immediately after solidifying to prevent drying.

Probes were mounted by the handle with ultraviolet-sensitive epoxy to a glass capillary and inserted into the agar at constant speed using a motorized micropositioner (Newport 860 Motorizer, Newport Corporation, Irvine, CA) controlled by custom firmware on an Arduino Uno with external 12-bit DAC (Adafruit Industries, New York City, NY). A camera (2 MP OEM USB microscope) was positioned stationary relative to the probe to image the insertion process. Insertion speed was varied from 20-240 μm/s in 20μm/s increments to observe the effect on whisker deflection. Probes were inserted into the agar a distance of 5-6 mm, then retracted 57% of the length of the arc radius. This retraction distance was twice the difference in vertical position between the tip of a fully folded whisker and a whisker splayed by an angle of *π*/2. The deflection of the top right whisker was quantified in each recorded video. The agar was repositioned between insertions to avoid repeatedly penetrating the same point.

Devices were qualitatively grouped upon initial observation into those that folded completely during insertion and those that did not. These are represented by filled and open circles, respectively, on the plots in Figure 1a-b. The devices that folded completely were *w* =3 μm, *r* =600, 800, 1000 μm, and *w* =4 μm, *r* =800, 1000 μm. Those five devices were each tested in agar *N* = 5 times for each of the twelve speeds.

In addition to the agar insertions, probe *w* = 4, *r* = 1000 μm was inserted into a 500μm-thick slice of porcine brain positioned between two microscope cover slips. The probe was imaged with infrared light given its reduced scattering through lipid-rich tissue compared with visible light. Because the mechanical properties of the brain slice were dependent on the force applied between the cover slips, this was not an accurate model of healthy cortical tissue, and these data were not quantified.

## IV. Results and Discussion

Operation of the splaying probes was initially validated visually by viewing live and recorded video of the devices’ insertion into 0.6% agar, simulating insertion into cortical tissue.

Qualitative observations suggested that those with a rotational compliance ≲20 rad/μN fold poorly during insertion and were not tested extensively; only data from the five device geometries listed in Section III-B are presented. These observations guided the estimation in Figure 1a that the equivalent point load exerted at the tip of each whisker was approximately 40 μN, with the caveat that the insertion drag force experienced by each probe was a function of the whisker length.

Figure 3 shows photographs of the resting position, folded position, and splayed position of whiskers of the probe variant *w* = 4, *r* = 1000 μm in both agar and porcine gray matter. Devices could be removed successfully from both agar and brain without damage to the whiskers or irreversible stiction between whiskers. As predicted in Figure 1b, the yield strain of the silicon whiskers was greater than the maximum strain experienced during normal operation. This is critical for a clinically viable device, as a neural recording array must not leave fragments in the brain when explanted. A drawback of this device, however, is that the extent of the tissue or agar damage during explantation is as wide as the probe with whiskers splayed.

**Fig. 3.**
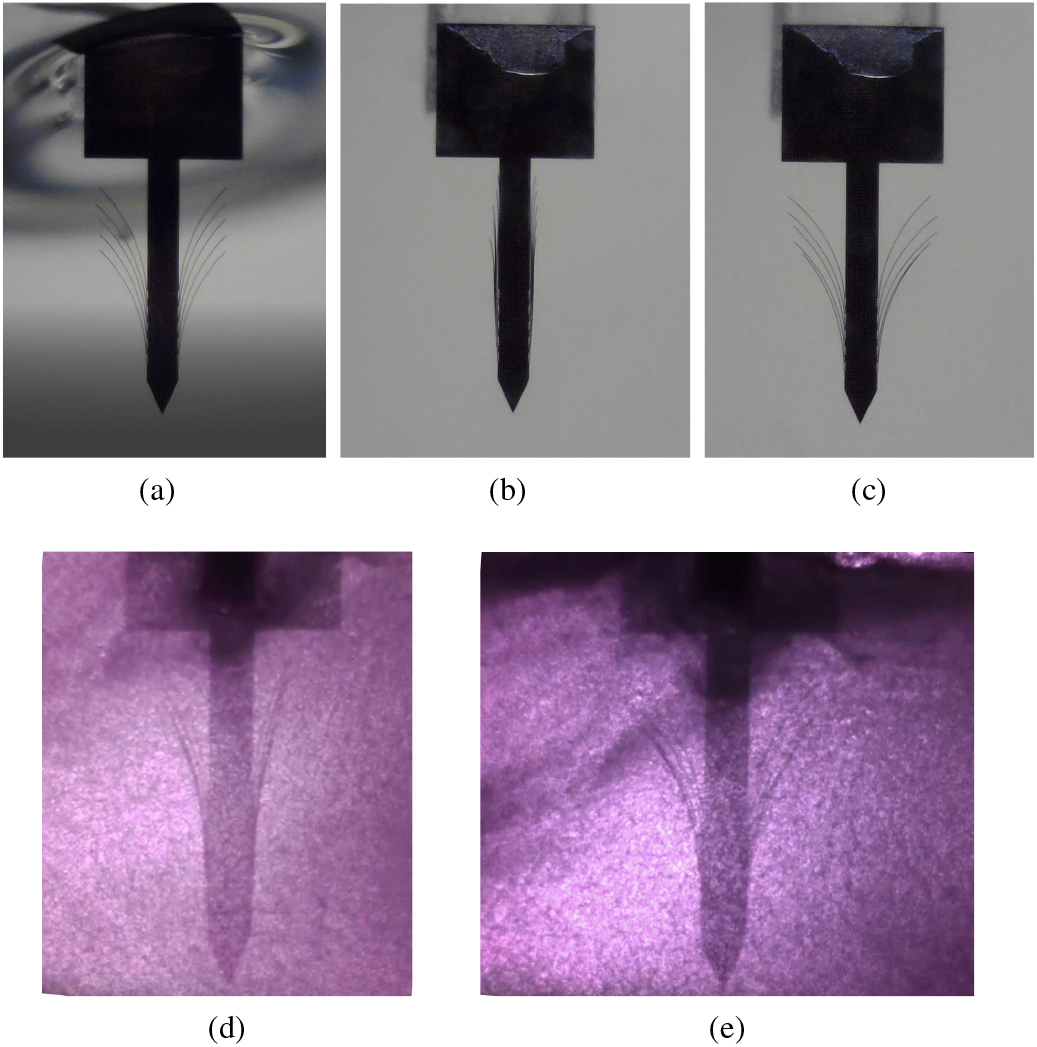
(a) Photograph of device before insertion (in deionized water), (b) during insertion into agar with whiskers folded, and (c) after retraction in agar with whiskers splayed. (d) photograph of probe during insertion into a porcine brain slice with whiskers incompletely folded, and (e) the same probe after retraction with the whiskers splayed relative to insertion. Scale: In all photographs, the probe body is 200 μm wide.

Results from the five insertions and retractions of probe *w* = 4, *r* =1000 μm at each of twelve speeds are summarized in Figure 4a,b. At all speeds, the whiskers folded nearly completely and splayed by at least 75 μm on average. Though the effect of insertion and retraction speed on deflection was rather weak, particularly given the large standard deviation of the data, a peak deflection was observed at 180 μm/s. This peak approximately corresponds to the maximum drag force as a function of insertions speed observed in literature for cortical tissue [13, 26, 27].

**Fig. 4.**
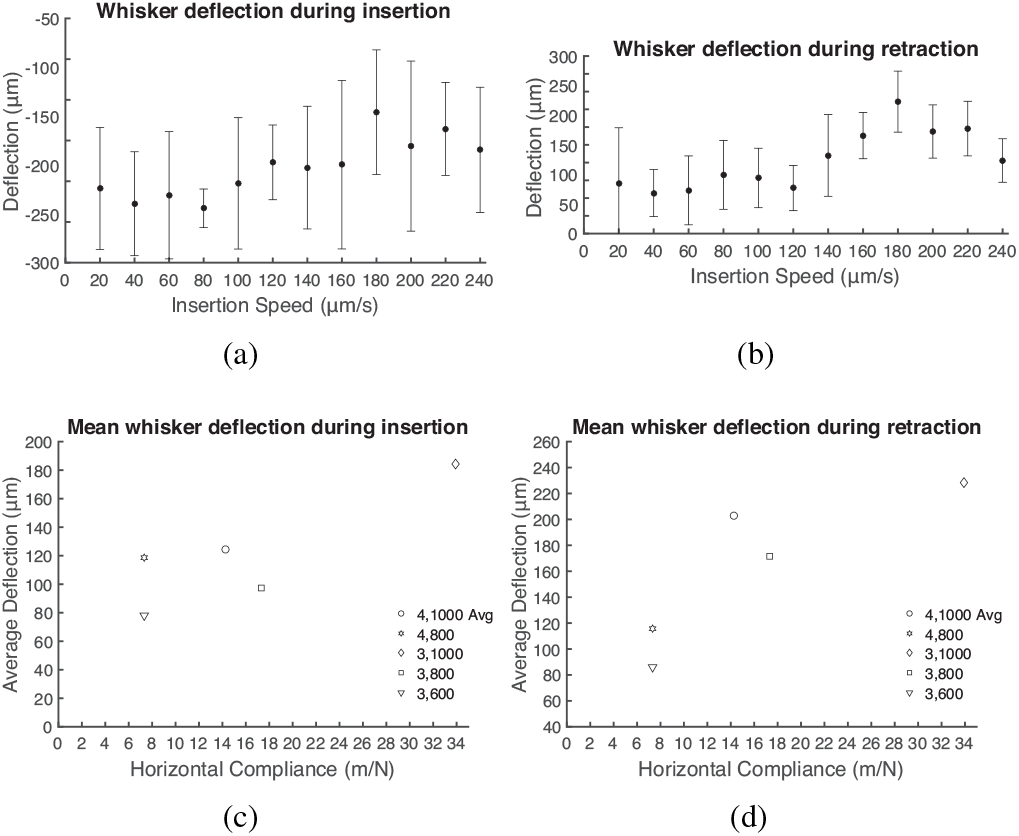
(a) Mean and standard deviation (N=5) of deflection versus speed during insertion for device *w* = 4, *r* =1000 μm. Deflection is measured as the magnitude of the difference in the x direction between the resting position of the top right whisker and its position at the end of insertion. (b) Mean and standard deviation (N=5) of deflection versus speed during retraction for device *w* = 4, *r* =1000 μm. Deflection is measured as the magnitude of the difference in the x-direction between the top right whisker’s position at the end of insertion and its final position after retraction. (c,d) Mean whisker deflection during insertion and retraction, averaged over all speeds, versus whisker compliance. Data are plotted for five device types, where values *w, r* in the legend correspond to beam width and arc radius in microns, respectively.

Deflection measurements from the five probe types tested were averaged over all speeds, and the resulting average deflection was plotted in Figure 4c,d against the compliance (inverse of the spring constant from Table II) of each of the probe geometries. As expected, the deflection was approximately linear with compliance. Differences in the degree of deflection for probes with the similar compliance values are explained by the greater insertion force experienced by probe geometries with longer whiskers. Additionally, we hypothesize that the design of the rectangular handle may have disrupted the agar during retraction, decreasing the degree of splaying observed. A tapered handle or longer probe body above the whiskers could mitigate this effect.

Testing in porcine neural tissue was limited to verifying that the splaying probes would still deflect. Because the mechanical properties of the tissue were dependent upon the force applied between the glass cover slips sandwiching the brain slice, quantitative data were not meaningful. However, initial qualitative testing suggested that the probes operated similarly in porcine brain slices compared with agar. The splayed position of the whiskers in Figure 3d,e was approximately 100 μm outside of the damage zone during insertion, suggesting that the adverse biological response near the tips of the whiskers would likely be minimal.

In addition to the designed function of the probe in placing the tips of the whiskers outside of the reach of the glial scar surrounding the probe body and tissue damage along the insertion track, prior literature suggests the whiskers can also aid in anchoring the probe and stabilizing the recording sites to micromotion [28, 29]. Because the whiskers are compliant relative to the cortical tissue, we expect that recording sites at the tips could track micromotion in the brain and remain stationary relative to target neurons while the probe body remains anchored to the skull.

A key limitation of this design is the low whisker density. Because the whiskers project from the anchors parallel to the probe body, their finite width imposes a minimum pitch (Figure 2b). Further, the design allows at most two recording sites at any cortical depth; this stands in contrast to conventional silicon probes that may have several recording sites across the width of the probe body at a given position along its length.

## V. Conclusion

This work has demonstrated a mechanical proof-of-concept neural probe in which fine whiskers deflect upon insertion and splay upon retraction to place the tips of the whiskers, on which future recording sites could be patterned, outside the typical reach of a glial scar that forms around such probe bodies. Because the whiskers’ cross-sectional dimensions are 3-5 μm, they are not expected to elicit an adverse biological response and thus may be amenable to stable chronic recording of individual units. The devices are monolithically fabricated in silicon, and the process is readily extensible to a variety of materials, including silicon carbide for device longevity [30].

Fabricating an electrically viable device requires only, at minimum, patterning conductive recording sites and traces and encapsulating the traces with an insulating material. Minor challenges anticipated in this translation involve managing deposited film stresses to avoid unwanted out-of-plane curling of the the whiskers. Compared to prior work in this space of actuated neural recording arrays, post-fabrication preparation of this device for surgical implantation is essentially identical to traditional shank-style silicon probes for neural recording.

Beyond the specific device presented here, this work advances the paradigm of incorporating traditional MEMS techniques of actuation and flexible components into the design of implantable devices.

## Acknowledgment

We thank the Berkeley Sensor and Actuator Center, the SwarmLab Research Centers, the Chan Zuckerberg Biohub, and the Center for Neural Engineering and Prostheses for their continued support.

## References

[1] Changhyun Kim and Kensall D Wise. “A 64-site multishank CMOS low-profile neural stimulating probe”. In: Solid-State Circuits, IEEE Journal of 31.9 (1996), pp. 1230–1238.

[2] Miguel AL Nicolelis et al. “Chronic, multisite, multielectrode recordings in macaque monkeys”. In: Proceedings of the National Academy of Sciences 100.19 (2003), pp. 11041–11046.

[3] Patrick K Campbell et al. “A silicon-based, three-dimensional neural interface: manufacturing processes for an intracortical electrode array”. In: Biomedical Engineering, IEEE Transactions on 38.8 (1991), pp. 758–768.

[4] Jozsef Csicsvari et al. “Massively parallel recording of unit and local field potentials with silicon-based electrodes”. In: Journal of neurophysiology 90.2 (2003), pp. 1314–1323.

[5] Jiangang Du et al. “Multiplexed, high density electrophysiology with nanofabricated neural probes”. In: PloS one 6.10 (2011), e26204.

[6] Jorg Scholvin et al. “Close-packed silicon micro-electrodes for scalable spatially oversampled neural recording”. In: Biomedical Engineering, IEEE Transactions on (2015), pp. 120–130.

[7] Carolina Mora Lopez et al. “An implantable 455-active-electrode 52-channel CMOS neural probe”. In: Solid-State Circuits, IEEE Journal of 49.1 (2014), pp. 248–261.

[8] Antal Berényi et al. “Large-scale, high-density (up to 512 channels) recording of local circuits in behaving animals”. In: Journal of neurophysiology 111.5 (2014), pp. 1132–1149.

[9] György Buzsáki. “Large-scale recording of neuronal ensembles”. In: Nature neuroscience 7.5 (2004), pp. 446–451.

[10] William M Reichert. Indwelling neural implants: strategies for contending with the in vivo environment. CRC Press, 2007.

[11] Roy Biran, Dave C Martin, and Patrick A Tresco. “The brain tissue response to implanted silicon microelectrode arrays is increased when the device is tethered to the skull”. In: Journal of Biomedical Materials Research Part A 82.1 (2007), pp. 169–178.

[12] John P Seymour and Daryl R Kipke. “Neural probe design for reduced tissue encapsulation in CNS”. In: Biomaterials 28.25 (2007), pp. 3594–3607.

[13] Vadim S Polikov, Patrick A Tresco, and William M Reichert. “Response of brain tissue to chronically implanted neural electrodes”. In: Journal of neuroscience methods 148.1 (2005), pp. 1–18.

[14] Paras R Patel et al. “Chronic in vivo stability assessment of carbon fiber microelectrode arrays”. In: Journal of neural engineering 13.6 (2016), p. 066002.

[15] Takashi D Yoshida Kozai et al. “Ultrasmall implantable composite microelectrodes with bioactive surfaces for chronic neural interfaces”. In: Nature materials 11.12 (2012), pp. 1065–1073.

[16] Paras R Patel et al. “Insertion of linear 8.4 *μ*m diameter 16 channel carbon fiber electrode arrays for single unit recordings”. In: Journal of neural engineering 12.4 (2015), p. 046009.

[17] Daniel Egert and Khalil Najafi. “New class of chronic recording multichannel neural probes with post-implant self-deployed satellite recording sites”. In: Solid-State Sensors, Actuators and Microsystems Conference (TRANSDUCERS), 2011 16th International. IEEE. 2011, pp. 958–961.

[18] Chong Xie et al. “Three-dimensional macroporous nanoelectronic networks as minimally invasive brain probes”. In: Nature materials 14.12 (2015), p. 1286.

[19] Daniel GD Egert. “Intracortical Neural Probes with Post-Implant Self-Deployed Electrodes for Improved Chronic Stability.” In: (2015).

[20] Stéphanie F Bernatchez, Patrick J Parks, and Donald F Gibbons. “Interaction of macrophages with fibrous materials in vitro”. In: Biomaterials 17.21 (1996), pp. 2077–2086.

[21] JE Sanders, CE Stiles, and CL Hayes. “Tissue response to single-polymer fibers of varying diameters: evaluation of fibrous encapsulation and macrophage density”. In: Journal of biomedical materials research 52.1 (2000), pp. 231–237.

[22] Warren Clarence Young and Richard Gordon Budynas. Roark’s formulas for stress and strain. Vol. 7. McGraw-Hill New York, 2002.

[23] Zhi-Jian Chen et al. “A realistic brain tissue phantom for intraparenchymal infusion studies”. In: Journal of neurosurgery 101.2 (2004), pp. 314–322.

[24] Farhana Pervin and Weinong W Chen. “Mechanically similar gel simulants for brain tissues”. In: Dynamic Behavior of Materials, Volume 1. Springer, 2011, pp. 9–13.

[25] Flavia Vitale et al. “Neural stimulation and recording with bidirectional, soft carbon nanotube fiber micro-electrodes”. In: ACS nano 9.4 (2015), pp. 4465–4474.

[26] Marleen Welkenhuysen et al. “Effect of insertion speed on tissue response and insertion mechanics of a chronically implanted silicon-based neural probe”. In: IEEE Transactions on Biomedical Engineering 58.11 (2011), pp. 3250–3259.

[27] CS Bjornsson et al. “Effects of insertion conditions on tissue strain and vascular damage during neuroprosthetic device insertion”. In: Journal of neural engineering 3.3 (2006), p. 196.

[28] A Cutrone et al. “A three-dimensional self-opening intraneural peripheral interface (SELINE)”. In: Journal of neural engineering 12.1 (2015), p. 016016.

[29] S Martel and T Fofonoff. “New approaches for the implementation of minimally invasive microelectrode arrays designed for brain-machine interfaces”. In: Engineering in Medicine and Biology Society, 2003. Proceedings of the 25th Annual International Conference of the IEEE. Vol. 4. IEEE. 2003, pp. 3794–3797.

[30] Camilo Diaz-Botia et al. “A silicon carbide electrode technology for the central and the peripheral nervous system”. In: Journal of Neural Engineering (2017).

